# Temporal dynamics of cytokine, leukocyte, and whole blood transcriptome profiles of pigs infected with African swine fever virus

**DOI:** 10.1101/2025.08.29.673161

**Authors:** Daniel W. Madden, Bianca Libanori Artiaga, Jessie D. Trujillo, Patricia Assato, Chester D. McDowell, Isaac Fitz, Taeyong Kwon, Konner Cool, Yonghai Li, Natasha N. Gaudreault, Igor Morozov, Juergen A. Richt

## Abstract

African swine fever virus (ASFV) is an important transboundary animal pathogen with significant impacts on the global swine industry. Overwhelming proinflammatory responses are a major virulence mechanism for ASFV, but the dynamics of these changes during clinical disease are not completely understood. We constructed a detailed portrait of the innate immune responses during acute African swine fever (ASF) at the cellular, transcriptomic, and cytokine levels. Samples serially obtained from infected piglets show progression of acute ASF is characterized by rapid increases in plasma type I interferons, TNF-α, IL-12p40, and IL-10, which coincide with the manifestation of clinical disease and viral DNAemia. Lymphocytes and natural killer (NK) cells progressively declined, with fluctuations in B cell, CD8+ T cell, and CD4+/CD8+ T cell populations. Blood monocytes and macrophages were highly variable throughout infection, with an abrupt spike in CD203+ mature macrophages immediately prior to death. Transcriptomic analysis of blood showed downregulation of cellular translation as early as 1 day post challenge (DPC), and significant upregulation of antiviral immune processes at 5 DPC and 7 DPC which overlapped with the onset of clinical disease. Together, these results present a highly detailed delineation of fatal ASF as involving an initial infection and damage of susceptible myeloid cells prior to symptomatic disease characterized by pro-inflammatory immune responses, lymphoid depletion, and clinical deterioration.

## 1. Introduction

African swine fever virus (ASFV) is a highly contagious pathogen that causes African swine fever (ASF), a lethal disease in domestic pigs and wild boar [1]. Following its introduction to the country of Georgia in 2007, virulent genotype II ASFV has spread throughout much of Europe and Asia, causing the deaths of millions of domestic pigs and resulting in billions of dollars in economic losses [2]. ASFV is readily transmissible between domestic pigs and circulates widely in Eurasian wild boar populations, complicating eradication efforts [3]. Most recently, ASFV was identified in the Dominican Republic, further highlighting the transboundary nature of this devastating disease [4].

Despite decades of research, progress toward development of an ASFV vaccine has remained difficult, in part due to the highly complex biology of the virus itself and its interaction with host animals. To date, two modified live vaccines (MLVs) for ASFV are commercially available in Vietnam, one of which was temporarily removed due to concerns regarding the safety, efficacy, and usage of the vaccine [5,6]. ASFV is a nucleocytoplasmic large DNA virus possessing a genome of 170-190kbp with over 150 open reading frames, and many of its encoded proteins have unknown or poorly defined functions [7–9]. Infection outcomes can vary widely depending on the virus strain and host susceptibility. Highly virulent ASFV isolates such as those currently circulating in Europe and Asia produce a hemorrhagic fever with mortality rates up to 100% in domestic pigs and Eurasian wild boar, while low-virulence strains produce subclinical or chronic infections in the same host species [10–12]. Wild African suid species, including warthogs (*Phacochoerus spp.*) and bush pigs (*Potamochoerus larvatus*), maintain the virus within endemic areas of southern and eastern Africa, but do not develop clinical disease after infection [3,13–15]. The high degree of variability among ASFV strains and infection outcomes complicates efforts to understand the factors governing ASFV protection and immunity.

Host immune responses to ASFV are highly complex. In susceptible pigs, virulent ASFV strains induce a massive pro-inflammatory immune response and the activation of monocytes-macrophages for which the virus has a tropism, and previous research has established this as a major mechanism of ASFV pathogenesis and lesion development [11,12,16–20]. Additionally, ASFV encodes an array of proteins with diverse immunomodulatory activities including inhibition of the induction and signaling of type I interferons (IFNs) and other cytokines, suppression of apoptosis, and decreasing antigen presentation [21,22]. The precise immunological mechanisms and correlates of protection associated with lethal vs. non-lethal ASF are not completely understood, hindering rational vaccine design and development.

Importantly, since highly virulent ASFV strains frequently produce death before robust adaptive immunity can be induced, innate and early adaptive immune responses play a significant part in ASFV immunopathology, and a more complete understanding of the myriad interactions between the virus and the immune system in acute fatal ASF is needed to guide vaccine development. To this end, this study serially assessed the progression of highly virulent genotype II ASFV infection in domestic pigs using several approaches including flow cytometry, cytokine quantification, and transcriptomic analysis to detail host immune responses throughout the acute disease process. By assessing these parameters simultaneously over the entire course of the disease, we have captured a comprehensive picture of the immune dysregulation that develops in domestic swine infected with virulent ASFV, which can be used to inform rational vaccine development efforts.

## 2. Materials and Methods

### 2.1. Cells and Viruses

ASFV Armenia 2007, a hemadsorbing genotype II strain isolated from domestic pigs, was obtained from the European Union Reference Laboratory for ASF (Centro de Investigación en Sanidad Animal, Instituto Nacional de Tecnología Agraria y Alimentaria [CISA-INIA]) [23]. Virus was propagated and titrated as 50% infectious dose in hemadsorbing units (HAD50/mL) on primary porcine alveolar macrophages (PAMs) as previously described by Carrascosa et al [24]. PAMs were obtained from healthy donor pigs via post-mortem bronchoalveolar lavage and were cultured in RPMI 1640 (Corning, Manassas, VA, USA) supplemented with 10% fetal bovine serum (FBS; Atlanta Biologicals, Flowery Branch, GA, USA), 1x MEM nonessential amino acids (Gibco, Waltham, MA, USA), and 1x antibiotic-antimycotic solution (Gibco). Cells were maintained in a humidified 37°C incubator with 5% CO2 atmosphere.

### 2.2. Animal experiments and clinical sampling

Six 9-week-old ASFV-naïve conventional outbred piglets were inoculated with 102 HAD50 of ASFV Armenia 2007 via intramuscular (IM) injection into the hindlimb. Clinical observations were performed daily, and animals were assigned total clinical scores for each day based on the sum of scores for 8 individual parameters including fever (0-4), body shape (0-3), liveliness (0- 3), respiratory signs (0-3), digestive signs (0-3), neurological signs (0-3), skin lesions (0-3), and ocular/nasal discharge (0-3), with a score of 0 indicating normal. Animals considered moribund or with a clinical score >10 were humanely euthanized by intravenous pentobarbital injection. Anticoagulated whole blood was collected by jugular venipuncture in EDTA and sodium heparin vacutainer tubes (BD, Franklin Lakes, NJ, USA) the day before challenge and at 1-, 3-, 5-, and 7- days post-challenge (DPC), and at the time of euthanasia. Whole blood for RNA isolation was collected in PAXgene blood RNA tubes (BD) the day before challenge and at 1-, 3-, 5-, and 7- DPC. Following euthanasia or expiration, all animals were necropsied for pathological evaluation. All work with potentially infectious materials was performed within high-containment BSL-3 and BSL-3Ag settings at the Biosecurity Research Institute at Kansas State University. Protocols were approved by the university’s Institutional Biosafety Committee (IBC; protocol #1314) and Institutional Animal Care and Use Committee (IACUC; protocol #4265).

### 2.3. Necropsy and gross pathological evaluation

All animals were necropsied following death or humane euthanasia for gross pathological evaluation and tissue collection. Gross pathological evaluation was performed by a veterinary pathologist as previously described [25,26]. Animals were assigned lesion severity scores for body condition (0-3), integument (0-3), cardiovascular system (0-6), liver (0-6), lung (0-9), kidney (0-3), spleen (0-6), GI tract (0-12), tonsil (0-6), mandibular lymph node (0-6), cranial mediastinal lymph node (0-6), renal lymph node (0-6), gastrohepatic lymph node (0-6), and prescapular lymph node (0-6), with a score of 0 indicating an absence of lesions. Disease severity for each animal was classified based on the sum of all individual lesion scores as mild (<30), moderate (31-47), or severe (>47).

### 2.4. Sample processing and storage

Sodium heparin blood tubes were centrifuged at 1,000 x g for 10 minutes to separate plasma and buffy coat. Plasma samples were then divided into single-use aliquots in Protein LoBind microcentrifuge tubes (Eppendorf, Hamburg, Germany) and immediately transferred to -80°C until use. Buffy coat obtained from plasma separation was then processed for white blood cell isolation and cryopreservation. Buffy coats were resuspended using phosphate-buffered saline (PBS; Gibco) with a volume equivalent to the amount of plasma removed from the sodium heparin blood tube. Then, 2mL of cell suspension was added to 15mL conical tubes containing 10mL/tube PBS. Tubes were gently mixed then centrifuged to pellet cells. Red blood cells were lysed using two treatments with ammonium chloride-based lysis buffer, and the remaining white blood cells were then washed with PBS and centrifuged to pellet. White blood cells were then resuspended in PBS + 2% FBS, filtered through 100µm cell strainers to remove debris, and total viable cell counts obtained using a Countess™ II Automated Cell Counter (Thermo Fisher Scientific, Waltham, MA, USA). Cells were then pelleted by centrifugation and resuspended in freezing medium composed of 90% FBS and 10% DMSO (Sigma-Aldrich, St. Louis, MO, USA), aliquoted into cryovials, placed in a controlled-rate cell freezing apparatus, and transferred to -80°C. Cells were stored in -80°C until thawed, fixed and stained for flow cytometry. PAXgene blood tubes were stored at -80°C until RNA purification was performed. Total RNA was purified from PAXgene blood RNA tubes using a PAXgene Blood miRNA kit (QIAGEN, Hilden, Germany) following the manufacturer’s protocol, and eluted RNA was stored at -80°C. EDTA whole blood was aliquoted into cryovials and stored at -80°C until use.

### 2.5. Viral DNA extractions and quantitative PCR (qPCR)

ASFV DNA concentration in whole blood was evaluated by quantitative real-time PCR (qPCR) targeting the ASFV p72 gene using TaqMan probes and primers developed by Zsak et al [27]. Viral DNA was obtained from EDTA whole blood by automated magnetic bead extraction as previously described [26,28]. PCR reactions were performed on a CFX96 Touch Real-Time PCR Detection system (Bio-Rad, Hercules, CA, USA) using PerfeCTa® FastMix® II as previously described [28], using nuclease-free water and plasmid encoding the target ASFV p72 sequence as positive and negative controls, respectively. Copy number (CN) of ASFV DNA was calculated using a standard curve derived from serial dilutions of positive control ASFV p72 plasmid and normalized per mL of blood (CN/mL).

### 2.6. Plasma cytokine quantification by ELISA

Cytokine levels in plasma samples were evaluated by sandwich ELISAs with analyte-specific antibody pairs performed on Nunc Maxisorp 96-well plates (Thermo Fisher Scientific). All ELISAs utilized biotinylated detection antibodies followed by incubation with streptavidin-HRP solution, color development with 3,3’,5,5’-Tetramethylbenzidine (TMB), and reaction termination with 450 nm stop solution. Assays for IL-1β, IL-10, IL-12p40, and TNF-α were performed using porcine DuoSet® ELISA kits (R&D Systems, Minneapolis, MN, USA), IFN-β was quantified using a swine IFN-β Do-It-Yourself ELISA kit (Kingfisher Biotech, St. Paul, MN, USA), and IL-2 and TGF-β1 were evaluated using commercial porcine matched antibody pair kits (Thermo Fisher Scientific), all following the manufacturer’s recommended protocols.

For evaluation of IL-1α concentrations, ELISA plates were first coated overnight with 200ng/well rabbit anti-pig IL-1α polyclonal antibody (Bio-Rad) diluted in 5x ELISA coating buffer (Bio-Rad). The following morning, plates were blocked with 3% bovine serum albumin (BSA; Sigma-Aldrich) in phosphate-buffered saline with 0.05% Tween-20 (PBST; Sigma-Aldrich) for two hours at room temperature. Plates were then washed with PBST and incubated with plasma samples diluted in PBST with 1% BSA for 90 minutes at room temperature. Plates were washed again and incubated with 100µL/well biotinylated rabbit anti-pig IL-1α detection antibody (Bio-Rad) diluted to 1.0µg/mL in PBST with 1% BSA for 1 hour at room temperature, followed by additional washes and incubation with 100µL/well streptavidin-HRP (R&D Systems) dilution for 1 hour. Plates were developed by adding 100µL/well 1-Step™ Ultra TMB- ELISA Substrate Solution (Thermo Fisher Scientific), and color development halted by addition of an equal volume of 450nm stop solution (Abcam, Cambridge, UK). Dilutions of recombinant pig IL-1α expressed in Pichia pastoris (Bio-Rad) were used for protein standards.

ELISA for detection of IFN-α was performed using mouse anti-pig IFN-α monoclonal antibody clones K9 (PBL Assay Science, Piscataway, NJ, USA) and F17 (Invitrogen, Waltham, MA, USA), with recombinant porcine IFN-α produced in yeast (Abcam) used for protein standard dilutions. Monoclonal antibody clone F17 was biotinylated using a LYNX Rapid Plus Biotin (Type 1) Antibody Conjugation Kit (Bio-Rad) following the manufacturer’s protocol. For the assay, monoclonal antibody clone K9 was diluted to 2.25µg/mL in 5x ELISA coating buffer and plates coated overnight at room temperature by adding 100µL/well coating antibody solution. The following morning, plates were washed with PBST and blocked for 2 hours at room temperature using a 3% BSA in PBST. Plasma samples and protein standards diluted in 1% BSA solution were added to the plates, which were then incubated for 90 minutes at room temperature, followed by PBST washes and addition of 100µL/well biotinylated clone F17 detection antibody diluted to 2.75µg/mL in 1% BSA solution. After incubating 1 hour, plates were washed and 100µL/well streptavidin-HRP dilution was added for 1 hour. Plates were then washed and developed with 100µL/well 1-Step™ Ultra TMB-ELISA Substrate Solution, followed by addition of an equal volume of 450nm stop solution.

To minimize the potential for analyte damage and degradation, all ELISAs were performed with single-use plasma aliquots which had not undergone any prior freeze-thaw cycles. All samples and standards were assayed in duplicate wells. For all ELISA plates, absorbance was read at 450nm using a BioTeK ELx808 plate reader (Agilent, Santa Clara, CA, USA). Blank-adjusted O.D. values for protein standard dilutions on each plate were used to generate sigmoidal four parameter logistic standard curves. Plasma concentrations for cytokines were determined by interpolating the blank-adjusted O.D. values for each sample into the appropriate standard curve and multiplying by the relevant dilution factor for each assay.

### 2.7. Flow cytometry

Cryopreserved white blood cells were thawed and washed with PBS, and cells were stained for flow cytometry as previously described [29]. Live/dead staining of cells was performed with LIVE/DEAD™ Fixable Green Dead Cell Stain Kit (Invitrogen) following the manufacturer’s instructions. Nonspecific antibody binding sites were blocked by incubating cells with recombinant rat IgG (Sigma-Aldrich), and cells were then stained for leukocyte surface markers using the antibodies outlined in Table 1. Three separate monoclonal antibody cocktails were used for cell staining. The first cocktail was used to distinguish B cells and T cells and consisted of antibodies targeting CD79a and CD3ε, respectively. The second cocktail contained antibodies targeting CD3ε, CD4, and CD8α for discrimination of T cell subsets and NK cells.

**Table 1.** Antibodies used for flow cytometry panels.

| Marker | Clone | Isotype | Source | Fluorochrome |
| --- | --- | --- | --- | --- |
| CD3ε | BB23-8E6-8C8 | Mouse IgG2aκ | BD Biosciences | PE-Cy7 |
| CD4 | 74-12-4 | Mouse IgG2aκ | BD Biosciences | PE |
| CD8α | 76-2-11 | Mouse IgG2aκ | Invitrogen | PE-Cy5 |
| CD14 | MIL2 | Mouse IgG2b | Bio-Rad | Qdot 800* |
| CD79a | HM47 | Mouse IgG1κ | Invitrogen | PE |
| CD163 | 2A10/11 | Mouse IgG1 | Bio-Rad | PE |
| CD203a | PM18-7 | Mouse IgG1 | Bio-Rad | NFB 660^ |

\*Conjugated in-house with Siteclick Qdot 800 Antibody Labeling Kit (Thermo Fisher Scientific)
^Conjugated in-house with NovaFluor™ Blue 660/120S Conjugation Kit (Thermo Fisher Scientific)

The third cocktail contained antibodies against CD14, CD163, and CD203a for characterization of monocytes and macrophages. Fluorescence-minus-one controls were included to confirm negative and positive populations. Stained cells were fixed with 4.2% formaldehyde solution then permeabilized and washed using a Cytofix/Cytoperm Fixation/Permeabilization Kit (BD) following the manufacturer’s protocol. Cell populations were analyzed using an Attune NxT flow cytometer (Thermo Fisher Scientific) equipped with a 488nm excitation laser. Collected raw data was analyzed using FlowJo version 10.7.0 software (Treestar, Palo Alto, CA, USA).

### 2.8. mRNAseq, differentially expressed genes, and gene ontology analysis

RNA extracted from PAXgene blood tubes was quantified and analyzed on a Qubit^TM^ 4 fluorometer (Thermo Fisher Scientific) using a Qubit^TM^ RNA Broad Range assay kit (Thermo Fisher Scientific) and a Qubit^TM^ RNA IQ assay kit (Thermo Fisher Scientific) following the manufacturer’s instructions and then normalized to 500ng. Library preparation was performed with Illumina Stranded mRNA Prep (Illumina, San Diego, CA, USA), with 16 samples per run. Briefly, mRNA was captured by beads, followed by a fragmentation reaction. mRNA was then used for first and second strand cDNA synthesis and adapter ligation. Dual indexing and library preparation was performed using IDT for Illumina DNA/RNA UD Indexes Set A (Illumina).

Library clean-up was done using AMPure XP beads (Beckman Coulter, Brea, CA, USA), followed by quantification using a Qubit^TM^ dsDNA Broad Range kit (Thermo Fisher Scientific). Libraries were normalized to 4 nM and denatured following the manufacturer’s instructions and run on the NextSeq 550 sequencer (Illumina) using a NextSeq 500/550 High Output Kit v2.5 (2x75 cycles) (Illumina). The loading concentration of the library was 1.8 pM, with 1% PhiX.

For analysis of host transcriptomics, reads generated from the sequencing runs were mapped to the Sus scrofa reference genome (Accession number: GCF_000003025.6) using Bowtie 2 [30]. Output SAM files from Bowtie 2 were sorted by QNAME using SAMtools [31] then fed into HTSeq [32] to generate gene counts based on the reference Sus scrofa genome. Counts were then analyzed with DEseq2 [33] to identify differentially expressed genes (DEGs) using an adjusted p-value cutoff of <0.05 as the threshold for significance. Lists of DEGs and their corresponding log2 fold-change from pre-challenge baseline abundance were made for each post-challenge timepoint and uploaded to PANTHER version 17.0 for gene ontology (GO) analysis [34]. Statistical enrichment tests were performed using the GO biological process annotation complete dataset (GO Ontology database. DOI: 10.5281/zenodo.7942786. Released 2023-05-10) to screen for host processes that were significantly up- or down-regulated based on the number of DEGs and changes in their relative abundance [35]. GO biological process annotations were sorted by false discovery rate (FDR)-adjusted p-value with a significance cutoff of p<0.05. For evaluation of ASFV transcriptomics, sequencing run reads were mapped to an annotated ASFV strain Armenia 2007 genome (unpublished, courtesy of David Meekins) and analyzed using a CLC genomics workbench v23.0.1 (QIAGEN) transcriptomics workflow. ASFV gene expression was measured as reads per kilobase of transcript per million reads mapped (RPKM).

### 2.9. Statistical analysis

Statistical analyses and graphing for body temperature, clinical scores, whole blood ASFV qPCR, plasma cytokine concentrations, and flow cytometry data were performed using GraphPad Prism v9.5.1 (GraphPad Software, Boston, MA, USA). Normality for datasets was assessed using the Shapiro-Wilk test. Data for body temperature, clinical scores, and whole blood ASFV qPCR was analyzed and compared to baseline pre-challenge values using the Kruskal-Wallis test with post-hoc Dunn’s multiple comparisons test. Plasma cytokine concentrations and flow cytometry results were analyzed using repeated measures one-way ANOVA with post-hoc Tukey’s multiple comparisons test for datasets with normally distributed values, or nonparametric Friedman’s test with post-hoc Dunn’s multiple comparisons test for datasets violating normality.

## 3. Results

### 3.1. Post-challenge clinical progression, virus qPCR, and pathology

Following IM challenge with Armenia 2007, piglets developed clinical disease characterized by rapidly progressing fever and symptoms consistent with acute ASF (Figure 1). All animals succumbed to ASF or met the criteria for humane euthanasia at 8 DPC – pigs #1, #2, #4, and #6 were humanely euthanized, while pigs #3 and #5 died of the disease. Pig #3 became febrile at 4 DPC, followed by pigs #2, #4, #5, and #6 at 5 DPC, and pig #1 at 6 DPC. Body temperatures for the group were significantly elevated from the pre-challenge baseline beginning at 6 DPC until the end of the study, and fever of ≥41.1 °C was observed for all animals immediately prior to death or euthanasia (Figure 1A). Clinical scores showed a similar pattern, becoming significantly elevated from pre-challenge baseline at 6 DPC and continuing to trend upward until the end of the study at 8 DPC; all animals had a clinical score ≥4 at their last recorded timepoint at 7 or 8 DPC (Figure 1B). ASFV DNA was detectable in blood beginning at 3 DPC for pig #3, and at 5 DPC for pigs #2, #4, #5, and #6. Pig #1 did not develop detectable levels of ASFV DNA until 7 DPC; consequently, a statistically significant elevation of the group mean viral DNA from baseline was first present at 7 DPC (Figure 1C).

**Figure 1.**
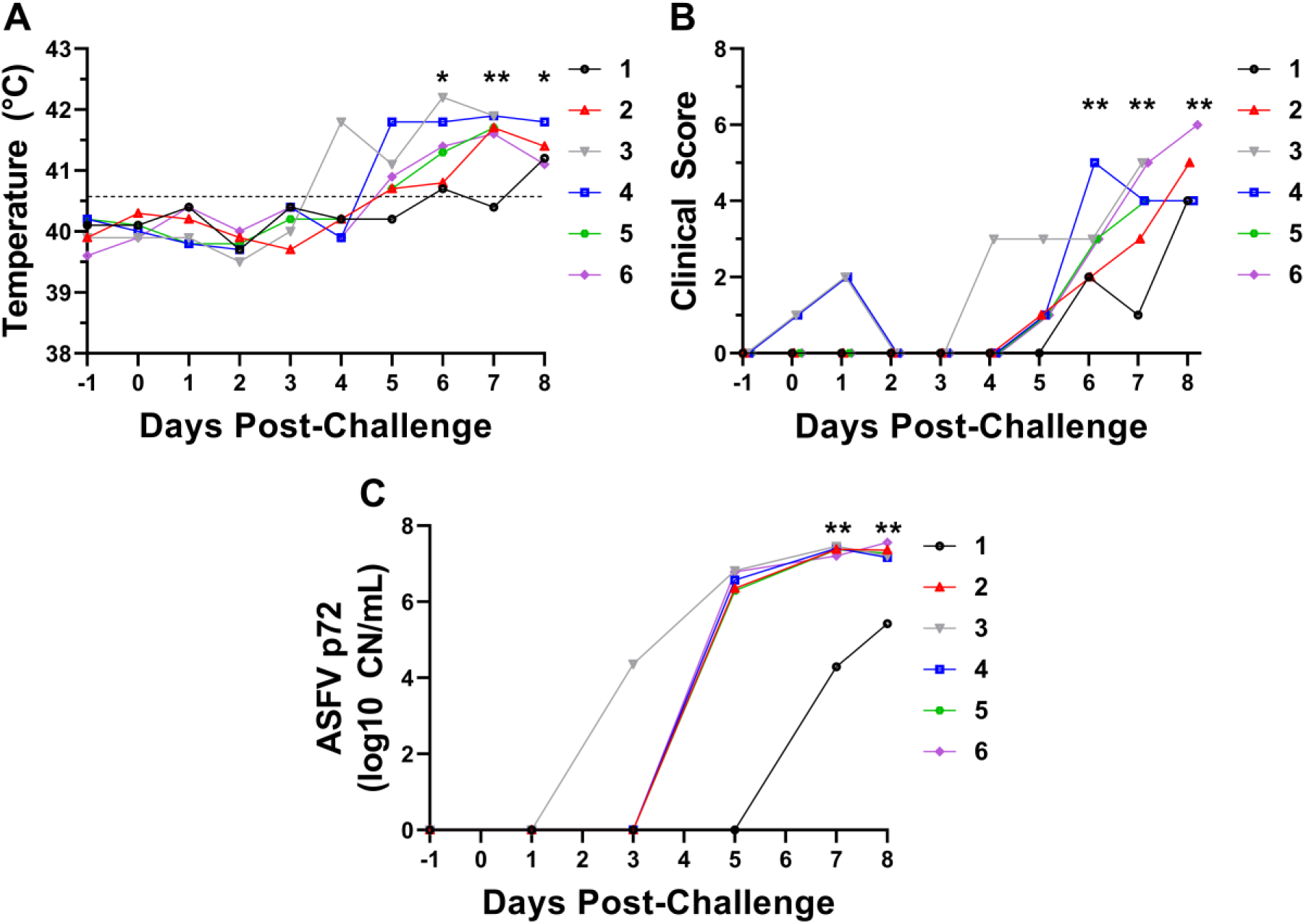
Body temperature (**A**), clinical scores (**B**), and ASFV p72 DNA levels (**C**). Statistical analysis was performed using the Kruskal-Wallis test with Dunn’s multiple comparisons to compare each timepoint to -1 DPC baseline. qPCR values are denoted as copy number of ASFV p72 gene per mL whole blood (CN/mL). *p<0.05, **p<0.01.

Animals challenged with Armenia 2007 developed gross pathological lesions consistent with acute ASF including decreased body condition, hemorrhagic lymphadenopathy and splenomegaly, pulmonary edema, consolidation pneumonia, and hepatopathy (Supplementary File S1). Pig #1 had the lowest total gross lesion score and was classified as having mild gross pathological lesions, consistent with clinical observations and PCR data showing this animal had the latest onset of fever and least amount of detectable viral DNA in the blood (Figure 1C, Supplementary File S1). Pigs #2, #4, #5, and #6 had total lesion scores classified as moderate disease, and pig #3 had a total lesion score classified as severe. Gross pathological abnormalities were noted for body condition, liver, lung, kidney, spleen, GI tract, mandibular lymph node, cranial mediastinal lymph node, renal lymph node, gastrohepatic lymph node, and prescapular lymph node in all animals. In addition, pig #3 had polyserositis and hemoabdomen (Supplementary File S1).

### 3.2. Plasma cytokine perturbations

A variety of pro- and anti-inflammatory cytokines showed altered expression levels in plasma over the course of virulent ASFV infection, including type I interferons, IL-10, IL-12p40, TNF-α, and TGF-β1 (Figure 2). IFN-α levels remained unchanged until 5 DPC when a significant spike in plasma concentrations was observed, which persisted to 7 DPC (Figure 2A). IFN-β levels increased significantly from 3 DPC to 7 DPC (Figure 2B). Concentrations of pro-inflammatory IL-1α and IL-1β were highly variable between animals prior to challenge and showed no significant changes from baseline at any timepoints (Figure 2C, D). IL-2 was undetectable in all animals prior to challenge, and highly variable responses were observed between animals post-challenge. Two pigs (#3 and #6) never developed detectable IL-2 levels, three pigs (#2, #4, #5) showed an initial increase in IL-2 at 1 DPC or 3 DPC which disappeared by 5 DPC and re-appeared at 7 DPC, and one pig (#1) had detectable levels of IL-2 only at the final 7 DPC timepoint; no statistically significant differences were detected between any timepoints (Figure 2E). No changes in IL-12p40 from baseline were seen until 7 DPC, when an abrupt 6-fold increase in the average plasma concentration was observed (Figure 2F). Similarly, TNF-α was stable throughout the first 6 days of the study until spiking significantly higher very late during infection at 7 DPC (Figure 2G). Levels of the anti-inflammatory cytokine IL-10 tended to increase early following ASFV infection, becoming significantly elevated from baseline at 5 DPC and peaking at 7 DPC (Figure 2H). In contrast, plasma TGF-β1, another cytokine with anti-inflammatory and immunoregulatory functions, did not significantly deviate from pre-challenge levels, and tended to progressively decrease from a peak mean at 1 DPC (Figure 2I).

**Figure 2.**
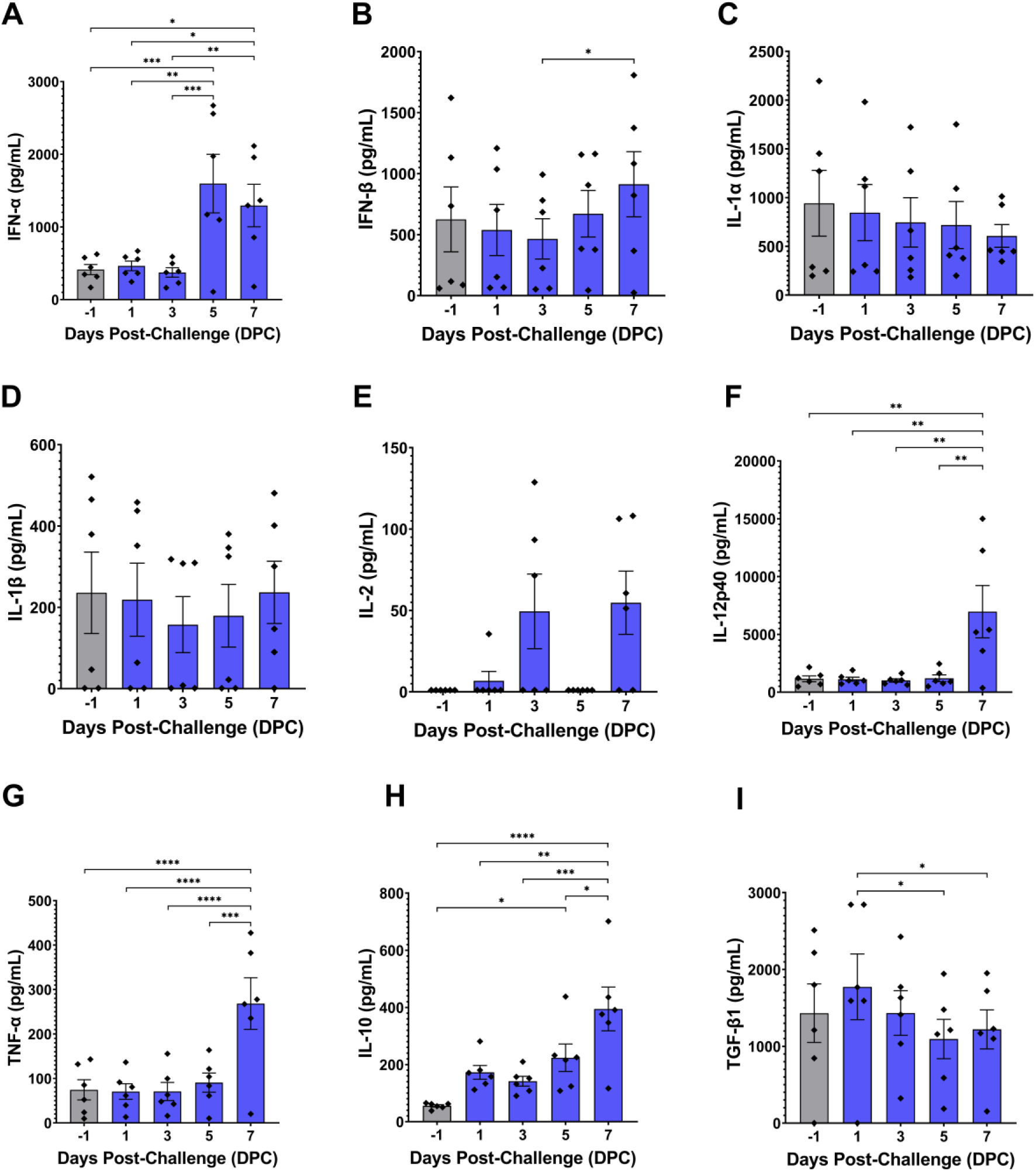
Plasma cytokine levels following ASFV challenge. Capture ELISAs were used to quantify levels of individual cytokines in plasma samples collected at the indicated timepoints. Normality of datasets was assessed with the Shapiro-Wilk test. (**A**), (**B**), (**C**), (**F**), (**G**), (**H**), and (**I**) were analyzed using repeated measures one-way ANOVA with post-hoc Tukey’s multiple comparisons test. (**D**) and (**E**) were analyzed using Friedman’s test with post-hoc Dunn’s multiple comparisons test. Error bars show mean with SEM (standard error of the mean). *p<0.05, **p<0.01, ***p<0.001, ****p<0.0001.

### 3.3. Changes in circulating leukocyte populations

Flow cytometry revealed changes in a variety of circulating leukocyte populations over the course of virulent ASFV infection, including total lymphocytes, CD8+ T cells, NK cells, B cells, monocytes, and macrophages. Total lymphocytes peaked at 1 DPC, followed by a steady decline over the course of ASFV infection. Total lymphocytes were significantly lower at 7 DPC compared to 1 DPC and 3 DPC in the T cell panel (Figure 3A) and B cells were significantly lower at 5 DPC and 7 DPC compared to earlier post-challenge timepoints (Figure 4A). The proportion of T cells remained relatively stable post-challenge, with no significant differences observed between any of the timepoints (Figure 3B, Figure 4B). CD4+/CD8α- helper T cells did not show any changes from baseline at any time point after infection (Figure 3C). However, CD8α+ T cell proportions changed over time. CD4-CD8α+ cytotoxic T cells were significantly lower at 5 DPC compared to their peak mean at 1 DPC (Figure 3D), while CD4+CD8α+ double- positive T cells were significantly elevated at 7 DPC compared to pre-challenge levels (Figure 3E). NK cells were initially stable during early infection but showed a significant decline at 5 DPC (Figure 3F). The percentage of B cells remained unchanged through 5 DPC but significantly decreased at 5 and 7 DPC (Figure 4C).

**Figure 3.**
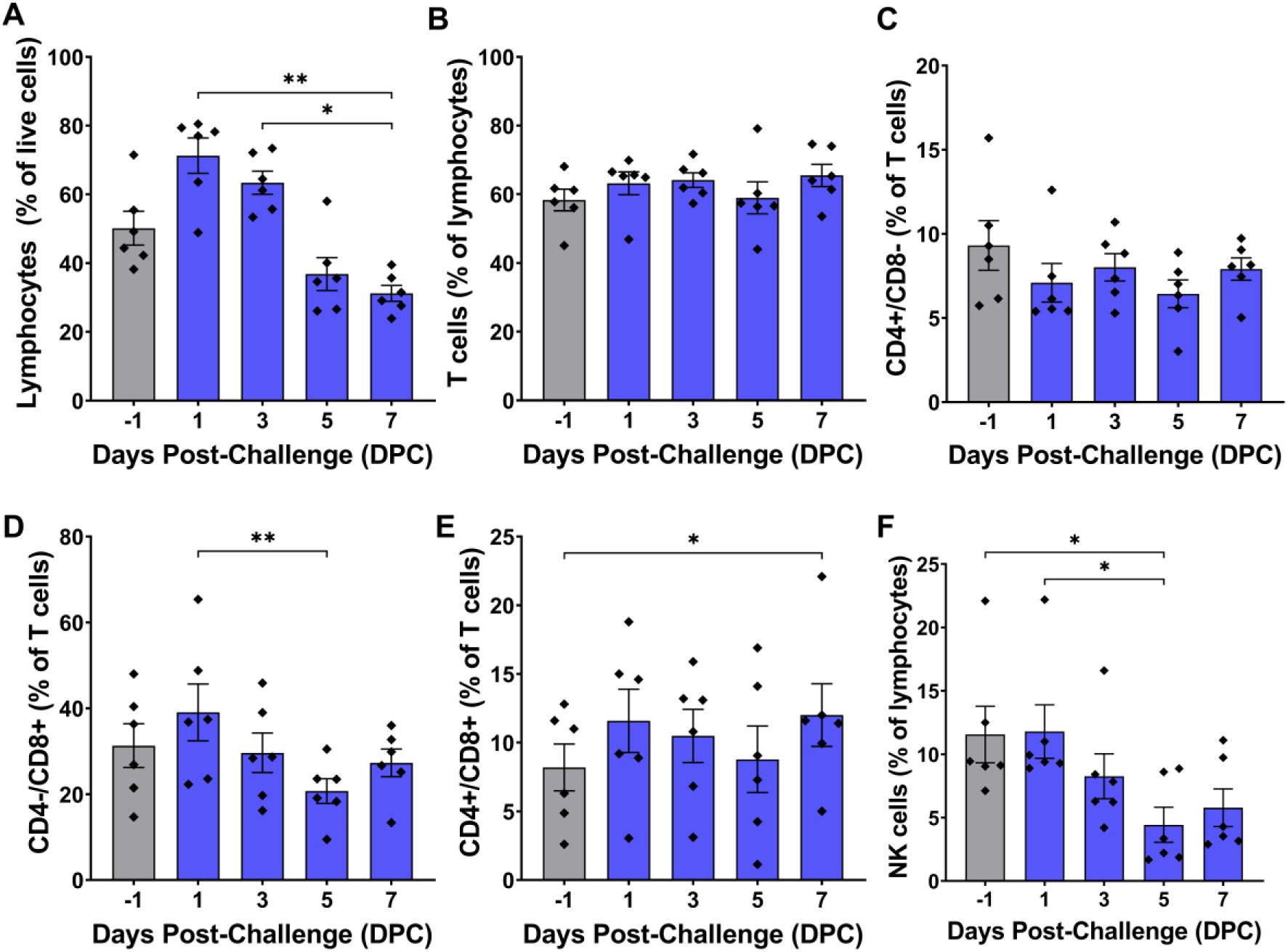
Circulating T cell subsets and NK cells during ASFV infection. White blood cells collected at each timepoint were fixed and stained using CD3ε, CD4, and CD8α antibodies listed in Table 1. T cells were defined as lymphocytes that were CD3ε+, while NK cells were defined as lymphocytes that were CD3ε-CD8α+. Individual cell phenotypes as a percentage of the total parent cell gate population are presented in each panel. Normality of cell populations was assessed with the Shapiro-Wilk test. (**A**), (**B**), (**C**), and (**F**) were analyzed using Friedman’s test with post-hoc Dunn’s multiple comparisons test. (**D**) and (**E**) were analyzed using repeated measures one-way ANOVA with post-hoc Tukey’s multiple comparisons test. Error bars show mean with SEM. *p<0.05, **p<0.01.

**Figure 4.**
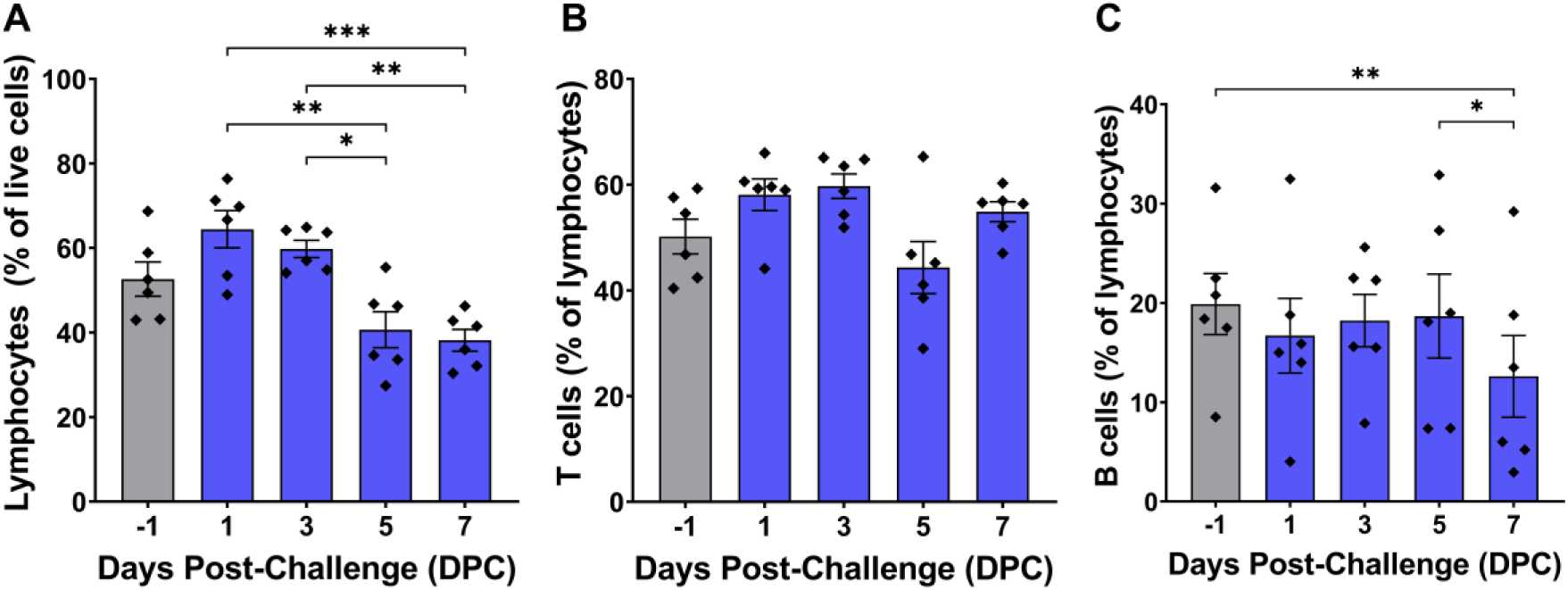
B cells during ASFV infection. White blood cells collected at each timepoint were fixed and stained using CD3ε and CD79a antibodies listed in Table 1. T cells were defined as CD3ε+ lymphocytes, while B cells were defined as CD79a+ lymphocytes. Individual cell phenotypes as a percentage of the total parent cell gate population are presented in each panel. Normality of cell populations was assessed with the Shapiro-Wilk test. (**A**) and (**C**) were analyzed using repeated measures one-way ANOVA with post-hoc Tukey’s multiple comparisons test. (**B**) was analyzed using Friedman’s test with post-hoc Dunn’s multiple comparisons test. Error bars show mean with SEM. *p<0.05, **p<0.01, ***p<0.001.

Monocytes and macrophages showed initial decreases in circulating populations very early at 1 DPC but subsequently rebounded to baseline levels at 3 DPC (Figure 5A, B). Monocytes continued to rise up to 5 DPC but dropped sharply by 7 DPC, just prior to death (Figure 5A). In contrast, total circulating macrophages declined to their lowest overall levels by 5 DPC but rebounded during the late stage of infection at 7 DPC (Figure 5B). Notably, CD203+ mature tissue macrophages remained relatively unchanged until a significant spike at 7 DPC when the population mean increased nearly 9-fold from baseline (Figure 5C).

**Figure 5.**
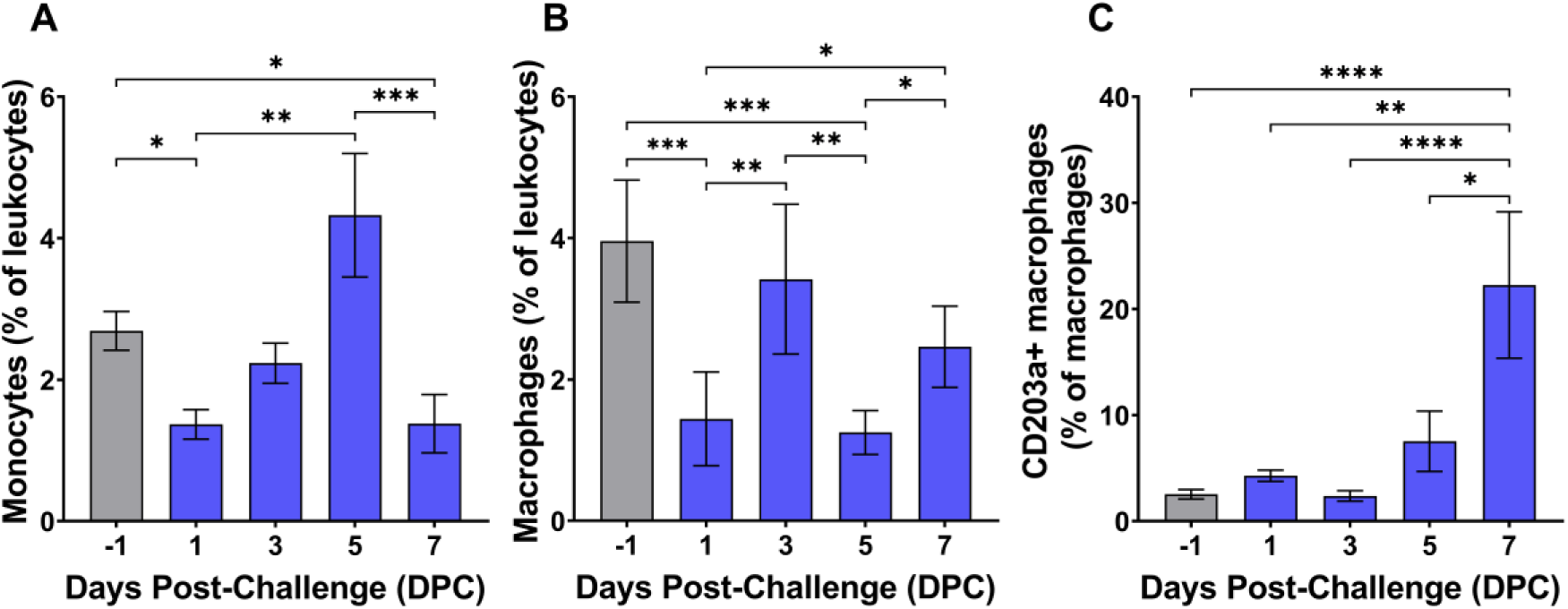
Monocytes (**A**), macrophages (**B**), and mature macrophages (C) during ASFV infection. Leukocytes collected at each timepoint were fixed and stained using CD14, CD163, and CD203a antibodies listed in Table 1. Monocytes were defined as CD14hiCD163+/-CD203a- while macrophages were classified as CD14loCD163+. CD203a+ macrophages were defined as mature tissue macrophages. Individual cell phenotypes as a percentage of the total parent cell gate population are presented in each panel. Normality of cell populations was assessed with the Shapiro-Wilk test. Data in all panels was analyzed using repeated measures one-way ANOVA with post-hoc Tukey’s multiple comparisons test. Error bars show mean with SEM. *p<0.05, **p<0.01, ***p<0.001, ****p<0.0001.

### 3.4. Differentially expressed genes (DEGs) and gene ontology (GO) processes

Host DEGs were detectable at various days post-challenge, with their number progressively increasing over the course of infection (Supplementary File S2). GO biological process analysis revealed downregulation of processes related to protein translation as well as amide and peptide metabolism as early as 1 DPC (Table 2). Consistent with this finding, 6 of the top 10 most downregulated DEGs by significance at 1 DPC encoded ribosomal proteins or peptidases (Supplementary File S2). There was no significant positive or negative enrichment from pre-challenge levels at 3 DPC. The largest number of DEGs was present at 5 DPC, when highly significant downregulation of processes relating to translation and peptide synthesis, amino acid metabolism, and ribosome function were evident, along with significant enrichment of processes relating to host defense and immune responses (Table 2). At 7 DPC, all enriched GO terms involved processes related to host immunity, regulation of viral replication, and stress responses (Table 2).

**Table 2.** Enriched GO biological processes during ASFV infection.

| DPC | GO Biological Process | ↑ or ↓ | # of DEGs | FDR p-value |
| --- | --- | --- | --- | --- |
| 1 DPC | amide biosynthetic process (GO:0043604) | ↓ | 17 | 4.68E-03 |
|  | amide metabolic process (GO:0043603) | ↓ | 18 | 5.82E-03 |
|  | peptide metabolic process (GO:0006518) | ↓ | 16 | 7.75E-03 |
|  | peptide biosynthetic process (GO:0043043) | ↓ | 16 | 1.16E-02 |
|  | translation (GO:0006412) | ↓ | 16 | 2.33E-02 |
|  | cytoplasmic translation (GO:0002181) | ↓ | 8 | 3.96E-02 |
| 5 DPC | peptide biosynthetic process (GO:0043043) | ↓ | 66 | 2.44E-09 |
|  | translation (GO:0006412) | ↓ | 66 | 3.66E-09 |
|  | peptide metabolic process (GO:0006518) | ↓ | 66 | 7.31E-09 |
|  | amide metabolic process (GO:0043603) | ↓ | 72 | 5.32E-08 |
|  | amide biosynthetic process (GO:0043604) | ↓ | 72 | 6.66E-08 |
|  | macromolecule biosynthetic process (GO:0009059) | ↓ | 80 | 3.79E-06 |
|  | gene expression (GO:0010467) | ↓ | 88 | 3.85E-06 |
|  | defense response to virus (GO:0051607) | ↑ | 30 | 4.13E-06 |
|  | defense response to symbiont (GO:0140546) | ↑ | 30 | 4.64E-06 |
|  | organonitrogen compound biosynthetic process (GO:1901566) | ↓ | 86 | 3.72E-05 |
|  | cellular nitrogen compound biosynthetic process (GO:0044271) | ↓ | 88 | 6.70E-05 |
|  | response to virus (GO:0009615) | ↑ | 34 | 1.47E-04 |
|  | ribonucleoprotein complex biogenesis (GO:0022613) | ↓ | 37 | 1.83E-04 |
|  | ribosome biogenesis (GO:0042254) | ↓ | 33 | 2.29E-04 |
|  | cytoplasmic translation (GO:0002181) | ↓ | 21 | 3.15E-04 |
|  | positive regulation of defense response (GO:0031349) | ↑ | 15 | 7.22E-04 |
|  | cellular biosynthetic process (GO:0044249) | ↓ | 94 | 7.40E-04 |
|  | regulation of innate immune response (GO:0045088) | ↑ | 18 | 1.48E-03 |
|  | regulation of defense response (GO:0031347) | ↑ | 23 | 1.76E-03 |
|  | negative regulation of viral process (GO:0048525) | ↑ | 17 | 2.04E-03 |
|  | positive regulation of innate immune response (GO:0045089) | ↑ | 13 | 2.18E-03 |
|  | regulation of response to biotic stimulus (GO:0002831) | ↑ | 21 | 3.45E-03 |
|  | non-membrane-bounded organelle assembly (GO:0140694) | ↓ | 20 | 3.48E-03 |
|  | organic substance biosynthetic process (GO:1901576) | ↓ | 101 | 4.15E-03 |
|  | regulation of viral process (GO:0050792) | ↑ | 22 | 5.49E-03 |
|  | biosynthetic process (GO:0009058) | ↓ | 103 | 5.62E-03 |
|  | negative regulation of viral genome replication (GO:0045071) | ↑ | 14 | 5.79E-03 |
|  | protein-RNA complex assembly (GO:0022618) | ↓ | 18 | 5.95E-03 |
|  | ribosome assembly (GO:0042255) | ↓ | 14 | 5.98E-03 |
|  | protein-RNA complex organization (GO:0071826) | ↓ | 18 | 6.17E-03 |
|  | ribosomal small subunit biogenesis (GO:0042274) | ↓ | 17 | 9.58E-03 |
|  | regulation of viral life cycle (GO:1903900) | ↑ | 21 | 9.73E-03 |
|  | cellular component biogenesis (GO:0044085) | ↓ | 76 | 1.13E-02 |
|  | positive regulation of response to biotic stimulus (GO:0002833) | ↑ | 14 | 1.23E-02 |
|  | cellular nitrogen compound metabolic process (GO:0034641) | ↓ | 121 | 1.24E-02 |
|  | ribosomal large subunit biogenesis (GO:0042273) | ↓ | 16 | 1.41E-02 |
|  | protein metabolic process (GO:0019538) | ↓ | 134 | 1.83E-02 |
|  | regulation of viral genome replication (GO:0045069) | ↑ | 16 | 2.01E-02 |
|  | defense response to other organism (GO:0098542) | ↑ | 54 | 2.36E-02 |
|  | regulation of response to external stimulus (GO:0032101) | ↑ | 33 | 2.92E-02 |
|  | organelle assembly (GO:0070925) | ↓ | 24 | 2.95E-02 |
|  | ribosomal small subunit assembly (GO:0000028) | ↓ | 8 | 2.98E-02 |
|  | protein-containing complex assembly (GO:0065003) | ↓ | 43 | 3.01E-02 |
|  | interspecies interaction between organisms (GO:0044419) | ↑ | 66 | 3.03E-02 |
|  | defense response (GO:0006952) | ↑ | 61 | 4.45E-02 |
| 7 DPC | defense response (GO:0006952) | ↑ | 81 | 8.12E-04 |
|  | interspecies interaction between organisms (GO:0044419) | ↑ | 80 | 8.68E-04 |
|  | defense response to virus (GO:0051607) | ↑ | 26 | 8.86E-04 |
|  | response to external biotic stimulus (GO:0043207) | ↑ | 78 | 9.32E-04 |
|  | response to other organism (GO:0051707) | ↑ | 78 | 1.07E-03 |
|  | defense response to symbiont (GO:0140546) | ↑ | 26 | 1.18E-03 |
|  | response to biotic stimulus (GO:0009607) | ↑ | 79 | 1.20E-03 |
|  | defense response to other organism (GO:0098542) | ↑ | 64 | 1.27E-03 |
|  | response to bacterium (GO:0009617) | ↑ | 24 | 7.90E-03 |
|  | response to virus (GO:0009615) | ↑ | 31 | 1.11E-02 |
|  | response to external stimulus (GO:0009605) | ↑ | 104 | 1.94E-02 |
|  | antimicrobial humoral response (GO:0019730) | ↑ | 8 | 2.06E-02 |
|  | regulation of defense response (GO:0031347) | ↑ | 45 | 2.35E-02 |
|  | positive regulation of defense response (GO:0031349) | ↑ | 27 | 2.61E-02 |
|  | response to stress (GO:0006950) | ↑ | 133 | 2.72E-02 |
|  | regulation of viral genome replication (GO:0045069) | ↑ | 15 | 3.64E-02 |
|  | negative regulation of viral genome replication (GO:0045071) | ↑ | 14 | 3.99E-02 |
|  | innate immune response (GO:0045087) | ↑ | 52 | 4.64E-02 |

Arrows indicate upregulated (↑) or downregulated (↓) processes. GO processes for each day are listed in descending order by level of significance. DPC = days post-challenge; GO = gene ontology; DEGs = differentially expressed genes; FDR = false discovery rate.

ASFV mRNAs were first detectable in whole blood at 5 DPC, when transcripts from 154 viral genes were identified, and by 7 DPC mRNAs from a total of 178 ASFV genes were observed (Supplementary File S3). The top 10 expressed ASFV genes for both timepoints are listed in Table 3, with 9 ASFV genes ranking among the most abundant in both timepoints. Members of MGF110 were highly represented post-challenge, with MGF110-7L the most abundantly transcribed ASFV gene at both 5 and 7 DPC. Additionally, other MGF110 members including MGF110-3L, MGF110-4L, and MGF110-5L-6L, were among the 10 most transcribed genes at both 5 and 7 DPC. Other highly expressed ASFV genes found at both timepoints include 285L, CP312R, E165R, K205R, and CP204L. K196R and I215L were the tenth most-transcribed viral genes identified at 5 DPC and 7 DPC, respectively.

**Table 3.** Top 10 ASFV genes by RPKM at 5 and 7 DPC.

|  | ASFV Gene | RPKM | Description | References |
| --- | --- | --- | --- | --- |
| 5 DPC | *MGF 110-7L\** | 1.17E+05 | Suppresses host cell translation | [82] |
|  | *285L\** | 1.16E+05 | Early viral gene product; membrane protein; nonessential | [89] |
|  | *CP312R\** | 9.57E+04 | ssDNA binding protein | [86] |
|  | *E165R\** | 9.06E+04 | dUTPase | [85] |
|  | *K205R\** | 8.20E+04 | Activates autophagy and NF-κB signaling | [91,92] |
|  | *MGF 110-3L\** | 7.91E+04 | Protein XP124L; virus morphogenesis | [80] |
|  | *CP204L\** | 6.70E+04 | Protein p30; virus internalization; endosome trafficking | [87,88,90] |
|  | *MGF 110-5L-6L\** | 5.58E+04 | Uncharacterized; nonessential | [83] |
|  | *MGF 110-4L\** | 3.74E+04 | Uncharacterized; has ER retention motif | [80] |
|  | *K196R* | 2.39E+04 | Thymidine kinase | [84] |
| 7 DPC | *MGF 110-7L\** | 1.45E+05 | Suppresses host cell translation | [82] |
|  | *K205R\** | 1.20E+05 | Activates autophagy and NF-κB signaling | [91,92] |
|  | *MGF 110-3L\** | 1.08E+05 | Protein XP124L; virus morphogenesis | [80] |
|  | *285L\** | 9.86E+04 | Early viral gene product; membrane protein; nonessential | [89] |
|  | *CP204L\** | 9.14E+04 | Protein p30; virus internalization; endosome trafficking | [87,88,90] |
|  | *E165R\** | 7.79E+04 | dUTPase | [85] |
|  | *MGF 110-5L-6L\** | 7.60E+04 | Uncharacterized; nonessential | [83] |
|  | *CP312R\** | 5.90E+04 | ssDNA binding protein | [86] |
|  | *MGF 110-4L\** | 4.61E+04 | Uncharacterized; has ER retention motif | [80] |
|  | *I215L* | 3.07E+04 | Ubiquitin-conjugating enzyme; inhibits type I IFN signaling | [93,94] |

Genes listed in both timepoints are denoted with \*. DPC = days post challenge. RPKM = reads per kilobase per million mapped reads.

## 4. Discussion

Immune dysregulation producing an overwhelming pro-inflammatory state is a major mechanism underlying ASFV pathogenesis and virulence [11,12,36]. However, a precise picture of the immunological mechanisms driving ASF progression remains elusive, hindered by the complexity of host-ASFV interactions, strain-dependent disease outcomes, inconsistencies between in vitro and in vivo findings, and variable experimental approaches. To overcome these limitations, we performed a virulent ASFV challenge experiment whereby a wide range of immune parameters were serially evaluated in the same group of piglets from baseline throughout the course of infection. This approach enabled the integration of data from diverse techniques - including clinical and post-mortem observation, molecular virological assays, cytokine quantification, flow cytometry of circulating immune cell populations, and transcriptomics - at defined timepoints post-challenge. Our results provide a detailed depiction of the evolving innate and early adaptive immune responses mounted by naïve piglets following infection with an epidemiologically relevant virulent genotype II ASFV strain.

All six piglets in this study developed clinical signs and post-mortem lesions characteristic of acute virulent ASF, though the development of symptoms and detectable viral DNA levels was slightly delayed in pig #1 compared to the other animals, and its gross necropsy findings were less severe (Figure 1, Supplementary File S1). Gross necropsy findings included widespread hemorrhagic lymphadenopathy, splenomegaly, hepatopathy, pulmonary edema and consolidation, and hemorrhage resulting from systemic vascular disease and disrupted hemostasis (Supplementary File S1). These lesions are highly consistent with descriptions of acute ASF in domestic pigs and wild boar [10,11].

Cytokines are a broad class of cell signaling proteins involved in the development and regulation of immune responses [37]. The widespread release of pro-inflammatory cytokines such as TNF-α from infected monocytes and macrophages seems to be a major mechanism governing ASFV pathogenesis [11,17–20,36]. ASFV also encodes a range of genes that modulate cytokine production and signaling, particularly type I interferons, to subvert host immunity. Several of these genes have been identified as key viral virulence factors [38]. Several studies have previously sought to evaluate the expression patterns of various pro- and anti-inflammatory cytokines during ASFV infection with the aim of characterizing the differences between host immune responses in lethal vs non-lethal ASF, sometimes producing inconsistent or conflicting results. For example, serum concentrations of the critical cytokines IFN-α and TNF-α have been independently reported to increase, decrease, or remain unchanged following virulent genotype II ASFV challenge in domestic pigs [39–42]. A variety of factors may explain these discrepancies, including variations in the *in vivo* behavior and virulence of different ASFV isolates as well as experimental design and methodology, including sampling timepoints.

The cytokines evaluated in this study regulate a diverse array of immunological functions including pro-inflammatory and anti-inflammatory responses, defenses against intracellular pathogens, T cell proliferation and differentiation, and the activity of innate and adaptive cytotoxic immune cells. They were selected based on prior research into ASFV-induced cytokine responses as well as their roles in mediating immune responses known to be important in the pathogenesis of ASF. Type I IFNs exert broad antiviral effects by inducing the expression of interferon-stimulated genes, and ASFV genes targeting type I IFN responses can mediate viral virulence in vivo [43–52]. IL-1α, which is constitutively expressed by epithelial and endothelial cells and released during cell necrosis, and IL-1β, which is primarily produced by monocytes and macrophages following activation of TLR signaling cascades, both mediate induction of multiple acute inflammatory processes including fever, prostaglandin secretion, histamine release, acute phase protein production, increased vascular permeability, and local infiltration of neutrophils and monocytes [53,54]. Previous studies have generally shown that infection with highly virulent ASFVs including E70 (genotype I, Europe), Armenia 2008 (genotype II, Caucasus), and SY18 (genotype II, China) can cause increases in serum IL-1α and IL-1β, while moderately virulent and attenuated ASFV strains do not [38,40–42,55–58]. TNF-α mediates a similar set of acute inflammatory responses and can also promote activation of macrophages and dendritic cells, induction of a pro-coagulant state in endothelial cells, and induction of cell death; it is produced primarily by macrophages and to a lesser extent by other immune cell types [59,60]. TNF-α is known to be an important mediator of ASFV pathogenesis in vivo, and multiple studies have demonstrated increases in serum TNF-α following challenge with moderately and highly virulent genotype I and II ASFV strains including E75 (genotype I; highly virulent), SY18 (genotype II, highly virulent), Armenia 2007 (genotype II, highly virulent) [16,40,41,57]; however, other studies have shown no significant changes in TNF-α following challenge with highly virulent SY18 [42,61]. IL-12 is produced by dendritic cells and other antigen presenting cells (APCs) and is important for cell-mediated immune responses through its ability to stimulate NK cell cytotoxicity, T cell differentiation into Th1 and cytotoxic T lymphocyte subsets, and IFN-γ production [62]. While cell-mediated immune responses seem to be essential for protection against lethal ASFV challenge, serum IL-12 levels during virulent ASFV infections remain poorly studied. However, SY18 infection has been reported to induce increasing IL-12 levels over the course of infection [41]. IL-2 is similarly critical for cell-mediated immunity via its role in stimulating T cell proliferation and differentiation; unfortunately, reports on the effects of highly virulent ASFV infection on serum IL-2 reveal inconsistent results [42,61,63]. Significant attention has been given to IL-10 responses during ASFV infection due to its role as an anti-inflammatory cytokine that regulates innate and adaptive immune responses by inhibiting antigen presentation and proinflammatory cytokine secretion by APCs, inhibiting Th1 responses, and promoting Treg differentiation [64,65]. In immunologically naïve animals, virulent ASFV challenge can produce increased serum IL-10 during late-stage infection immediately prior to death, though this is inconsistently observed [41,42,61]. However, vaccination/challenge studies using attenuated ASFV strains for immunization followed by virulent virus challenge tend to show a negative correlation between IL-10 secretion and survival following lethal challenge, and an association between IL-10 levels and an absence of long-term immunity [38,66–68]. TGF-β1, another major anti-inflammatory mediator capable of counteracting pathological immune responses, has been less extensively investigated, but appears to remain stable during infection with ASF strains Netherlands86 (genotype I, Europe) and SY18 (genotype II, China) ASFVs [41,57,69].

Our results show that virulent genotype II ASFV infection induces production of pro-inflammatory cytokines including type I interferons, IL-12p40, and TNF-α (Figure 2). IFN-α was the major type I interferon induced by infection, with plasma concentrations becoming significantly elevated at 5 DPC and 7 DPC (Figure 2A). In contrast, IFN-β concentrations tended to remain more stable with no significant difference from baseline observed at any timepoint post infection, though levels at 7 DPC were significantly higher than at 3 DPC (Figure 2B).

Significant changes in IL-12p40 and TNF-α were not observed until 7 DPC (Figure 2F, G). The late increases in these pro-inflammatory cytokines coincided with the development of significant elevations in body temperature and severe clinical signs at 6 DPC and occurred after the majority of animals developed detectable levels of ASFV DNA in the blood (Figure 1). While no significant changes were seen in the concentrations of IL-1α or IL-1β, there was a large amount of inter-animal variability with the levels of these cytokines which could have masked subtle changes (Figure 2C, D). Levels of IL-2 were below the limit of detection for all animals prior to challenge, and while 4/6 animals had detectable IL-2 at 7 DPC, no significant difference was noted (Figure 2E). This is consistent with the observed lack of significant changes in total T cell proportions at all post-challenge timepoints (Figure 3B). Interestingly, opposing responses were seen with the anti-inflammatory cytokines IL-10 and TGF-β1. Plasma IL-10 progressively increased over the course of infection and peaked immediately prior to death, while TGF-β1 was significantly decreased at 5 DPC and 7 DPC compared to its peak at 1 DPC (Figure 3H, I).

Multiple previous challenge studies have shown an association between increases in IL-10 and decreased survival following ASFV infection, suggesting IL-10 production during mid or late-stage infection may dampen protective immune responses and be detrimental to host survival [66–68]. The lack of TGF-β1 induction seem to support the detrimental effects of IL-10 induction.

Flow cytometry on PBMCs revealed changes consistent with both stimulation and aberrant adaptive immune responses. Populations of monocytes and macrophages, the primary cellular targets of ASFV infection, were highly variable following challenge and began fluctuating very early in the course of infection. Both cell types showed an initial significant decrease at 1 DPC followed by recovery at 3 DPC. Subsequently, monocyte populations remained relatively stable until sharply declining at 7 DPC, while macrophage populations dipped at 5 DPC but recovered immediately prior to death (Figure 5A, B). Interestingly, the proportion of CD203a+ macrophages abruptly spiked at 7 DPC to an average level nearly 9-fold higher than was seen prior to challenge (Figure 5C). CD203a, also known as SWC9, is a marker for differentiation of blood monocytes into macrophages, and expression of this molecule has been associated with increased cellular susceptibility to ASFV infection in vitro [70,71]. The human ortholog of CD203a, also known as ectonucleotide pyrophosphatase phosphodiesterase 1 (ENPP1), functions as an important negative regulator of type I interferon induction via the cGAS-STING signaling pathway by regulating extracellular cGAMP [72]. ENPP1 expression has been shown to facilitate pseudorabies virus infection in porcine cells by reducing IFN-β secretion, and virulent ASFV inhibits cGAS-STING signaling though multiple mechanisms [49–52,73]. This suggests that CD203a expression on macrophages could be an important regulator of type I interferon induction during ASFV replication in macrophages. Further evaluation of CD203a+ macrophages during in vivo ASFV infection is, therefore, warranted.

In contrast to myeloid-derived monocytes/macrophages, lymphoid cell populations displayed an initial trend toward increasing lymphocyte proportions at 1 DPC, followed by a progressive decline during later time points after infection that was mainly attributable to decreases in NK and B cell populations. The percentage of T cells remained relatively stable over time, while NK and B cells showed significant decreases at 5 and 7 DPC, respectively (Figure 3, Figure 4). CD4+/CD8α- helper T cells did not vary significantly throughout infection, while CD4-/CD8α+ cytotoxic T lymphocytes (CTLs) decreased from 1 DPC to 5 DPC (Figure 3C, D). In contrast, CD4+/CD8α+ double positive T cells were significantly elevated at 7 DPC (Figure 3E). CD8α+ cells are known to be important mediators of ASFV immunity, and depletion of these cells abolishes vaccine-induced protection against virulent ASFV challenge [74]. Defective CD8α+ T cell responses characterized by decreased perforin expression have previously been reported after virulent ASFV infection in domestic pigs and wild boar [75]. While we did not analyze the functional capacity of T cell subpopulations in this study, the observed lack of increases in helper T cell and CTL populations coupled with poor IL-2 induction is suggestive of a defective adaptive immune response.

Following ASFV inoculation, changes in the host transcriptome were detectable in whole blood as early as 1 DPC (Table 2; Supplementary File S2). The earliest detectable changes in GO biological processes were related to negative enrichment of genes involved biological processes related to protein translation and amino acid and peptide metabolism (Table 2). Processes relating to translation and ribosome assembly were similarly downregulated at 5 DPC, albeit to a more significant degree (Table 2). ASFV has previously been demonstrated to modulate cellular amino acid metabolism within infected PAMs in vitro, with early-stage infection producing increases in multiple amino acids including aspartate, glutamate, and arginine as well as enrichment of gene pathways relating to amino acid biosynthesis, and late-stage infection showing depletion of cellular amino acids [76]. The discrepancy between those in vitro results and the in vivo data presented here could be explained by multiple factors including the high MOI (1) used by Xue et al [76] to infect macrophages and fundamental differences in stress responses exhibited by cells in culture vs cells circulating in a dynamic in vivo system.

Positive enrichment of metabolic processes relating to innate and antiviral immune responses appeared at 5 DPC, and by 7 DPC the only significant changes seen were related to positive enrichment of immune response pathways (Table 2). These changes in immune-related processes overlap temporally with the appearance of significant changes in plasma cytokine levels (Figure 2) as well as the earliest detectable ASFV transcripts (Table 3). These results broadly agree with a previous study evaluating DEGs in whole blood samples collected at 7-10 DPC from pigs challenged with ASFV Georgia 2007, a strain very closely related to the Armenia 2007 strain used in this study, which showed enrichment of genes involved in RIG-I-like receptor signaling, an important antiviral innate immune pathway [77]. Four of the 20 DEGs showing the highest log2 fold changes in that study (S100A8, S100A9, LTF, and SIGLEC1) are among the top 20 DEGs identified at 7 DPC in the present study (Supplementary File S2).

Enrichment of innate immune responses in the organs of experimentally infected pigs following challenge with a virulent genotype II ASFV isolate from Vietnam has also been reported, with some pathways showing enrichment as early as 1 DPC; the earlier appearance of those reported changes compared to the present study likely reflects differences between samples being tested (organ tissue vs whole blood) and challenge inoculum dose (107 vs 102 HAD50) [78]. Overall, the host transcriptomic data presented here are consistent with immune response and metabolic changes known to occur during ASFV infection and provide valuable information on the temporal dynamics of these changes during disease progression.

ASFV gene transcripts were detectable at 5 DPC and 7 DPC, and 9 of the 10 most abundantly expressed viral genes were shared between both timepoints (Table 3, Supplementary File S3). MGF110-7L was the most abundantly transcribed gene at both timepoints, and other MGF110 genes, including -3L, -4L, and -5L-6L were also highly represented at both days. MGF110 members have previously been reported to be among the most transcribed ASFV genes in whole blood from Georgia 2007-infected pigs prior to death [77]. Unfortunately, the biological functions of many MGF110 genes are poorly characterized. These genes are transcribed beginning early after infection and exhibit a significant degree of amino acid conservation within the central domains of the predicted proteins [79,80]. At least two members of this family, MGF110-3L (also known as XP124L) and -4L, possess C-terminal endoplasmic reticulum (ER) retention motifs, and MGF110-4L is likely involved in incorporation of ER membrane components into maturing virions during viral morphogenesis [80,81]. MGF110-7L suppresses host cell translation and is involved in cellular stress granule formation, though it is unknown whether this is important for ASFV virulence [82]. The function of MGF110-5L-6L is not known, but this gene is nonessential for ASFV Georgia 2007 virulence in vivo [83]. Multiple ASFV proteins involved in DNA processing and nucleotide metabolism, including ssDNA binding protein CP312R, dUTPase E165R, and thymidine kinase K196R, showed high levels of early mRNA transcription which seems indicative of active viral genome replication [84–86].

ASFV genes C204L and 285L both encode membrane proteins. While C204L encodes p30, an ASFV protein known to be highly immunogenic and involved in virus internalization and endosome trafficking, the gene product of 285L is not well characterized, although it is known to be nonessential for in vitro virus replication [87–90]. Immunomodulatory viral genes were also among the most highly transcribed. K205R, which stimulates autophagy and NF-κB signaling, and inhibits type I IFN signaling through its interaction with IFNAR1 and IFNAR2, was the second most abundantly transcribed ASFV gene by 7 DPC [91,92]. Similarly, I215 inhibits type I IFN induction and signaling by promoting degradation of cellular IRF9 and STAT2 [93,94].

Taken together, these results highlight the temporal progression of virulent ASFV infection in domestic pigs, consisting of two distinct phases: (i) an initial subclinical infection characterized by perturbations in monocyte/macrophage populations and alterations in host metabolic and biosynthetic processes, along with restricted virus replication and global immune stimulation, and (ii) a later stage involving fulminant inflammatory responses and virus dissemination in the animal. Follow-up investigations of these parameters in well-defined ASFV-infected cell populations isolated through FACS analysis, single cell RNAseq or spatial transcriptomics could provide additional details into the temporo-spatial dynamics of ASFV infections and host responses. The significant increase in CD203a+ macrophages during the terminal phase of acute ASF warrants further investigation into the role of this marker during ASFV infection progression in vitro and in vivo. Additionally, the abundant transcription of different MGF110 genes at 5 DPC and 7 DPC highlights the need for additional research into the biological functions of these gene products.

## Supplementary Materials

Supplementary file S1: gross pathology scoring; Supplementary file S2: porcine DEGs identified by RNAseq; Supplementary file S3: ASFV genes identified by RNAseq.

## Author Contributions

Conceptualization, D.W.M. and B.L.A.; methodology, D.W.M., B.L.A., and J.D.T.; data acquisition, D.W.M., B.L.A., P.A., J.D.T., C.D.M., I.F., T.K., K.C., Y.L., N.G., I.M., and J.A.R.; data analysis, D.W.M., B.L.A., and P.A.; writing —original draft preparation, D.W.M.; writing— review and editing, B.L.A., P.A., J.D.T., C.D.M., I.F., T.K., K.C., Y.L., N.G., I.M., and J.A.R.; project administration, J.A.R.; funding acquisition, J.A.R. All authors have read and agreed to the published version of the manuscript.

## Institutional Review

The animal study protocol was approved by the Kansas State University Institutional Biosafety Committee (IBC; protocol #1314, approved 09/18/2018) and Institutional Animal Care and Use Committee (IACUC; protocol #4265.5, approved 05/21/2021).

## Acknowledgements

The authors would like to thank Carmina Gallardo and the European Union Reference Laboratory for African Swine Fever for providing the virus isolate used for this study.

